# A highly multiplexed melt-curve assay for detecting the most prevalent carbapenemase, ESBL and AmpC genes

**DOI:** 10.1101/842963

**Authors:** T. Edwards, C. Williams, Y. Teethaisong, J. Sealey, S. Sasaki, G. Hobbs, L.E. Cuevas, K. Evans, E.R. Adams

**Author notes:** Correspondence to TE.

## Abstract

Resistance to third generation cephalosporins and carbapenems in Gram-negative bacteria is chiefly mediated by beta-lactamases including ESBL, AmpC and carbapenemase enzymes. Routine phenotypic detection methods do not provide timely results, and there is a lack of comprehensive molecular panels covering all important markers.

An ESBL/carbapenemase HRM assay (SHV, TEM, CTX-M ESBLs, and NDM, IMP, KPC, VIM and OXA-48-like carbapenemases) and an AmpC HRM assay (16S rDNA control, FOX, MOX, ACC, EBC, CIT and DHA) were designed, and evaluated on 111 Gram-negative isolates with mixed resistance patterns.

The sensitivity for carbapenemase, ESBL and AmpC genes was 96.7% (95%CI:82.8-99.9%), 93.6% (95%CI:85.7-97.9%) and 93.8% (95%CI:82.8-98.7%), respectively with a specificity of 100% (95%CI:95.6-100%), 93.9% (95%CI:79.8-99.3%) and 93.7% (95%CI:84.5-98.2%).

The HRM assays enable the simultaneous detection of the fourteen most important ESBL, carbapenemase and AmpC genes and could be used as a molecular surveillance tool or to hasten detection of AMR for treatment management.

## 1. Introduction

The widespread resistance to first- and second-line antibiotics is one of the greatest challenges for the management of patients infected with Gram-negative bacteria, such as *Escherichia coli* and *Klebsiella pneumoniae* ^1^. The spread of Antimicrobial resistance (AMR) is facilitated by the organisms ability to maintain and transmit plasmids and other mobile elements containing resistance genes, such as those encoding beta-lactamases ^2^.

Resistance to 3^rd^ generation cephalosporins is frequently caused by extended spectrum beta-lactamase (ESBL) or AmpC genes, which are frequently mediated by plasmids ^3^. There are three main classes of ESBL; the TEM, SHV and CTX-M types, with the latter being the most prevalent class in the United Kingdom, and worldwide ^4,5^.

Infection with ESBL producers increases the length of hospitalization and mortality ^6^ and accounts for approximately 10% of *E. coli* isolates from serious blood stream infections in the UK ^5,7^. Reports from global surveillance networks have shown that ~15% of *E. coli* isolates in hospitals are ESBL producers, with a greater prevalence ranging from 20% to 60% in China and Southeastern Asian countries ^8^. AmpC genes are chromosomally carried by some members of the *Enterobacteriaceae*, including *Enterobacter spp* and *Citrobacter freundii complex*, and confer resistance to third generation cephalosporins and cephamycin, and plasmid mediated AmpC genes have enabled their spread throughout the *Enterobacteriaceae* ^9^. Acquired AmpC production in the Enterobacteriaceae is less frequently encountered than ESBL production, but is increasing ^8^, notably in Southeast Asia where 12% of *E. coli* in hospitals have plasmid-mediated AmpC genes ^10^.

Resistance to carbapenems, which are used as second line drugs for cephalosporin resistance, is becoming widespread. Carbapenemases include the enzymes VIM, IMP and NDM, KPC, and OXA carbapenemases ^11^. The prevalence of carbapenemase producers varies wildly worldwide, with <1% of invasive *K. pneumoniae* in the UK carrying these genes, compared with 66.9% in Turkey ^12^. The treatment options for infections of carbapenemase-producing organisms are limited and associated with poor clinical outcome ^13^, a 2-fold increase in mortality ^14^ and higher treatment costs ^15^. Determining antimicrobial susceptibility is a key diagnostic component to increase the likelihood of treatment success and to enable antimicrobial stewardship ^16^.

The detection of resistance to third generation cephalosporins and carbapenems is usually established by phenotypic tests such as the CLSI double disk potentiation method ^17^, the Etest (Biomerieux, France) 18 and the modified Hodge test ^19^. These methods provide key information on drug susceptibility phenotype, but require two 24-hour incubation periods, one to isolate the organism and a further incubation to establish the drug sensitivity profile. These steps usually take 48-96 hours with antibiotic prescription initiated empirically, resulting in the unnecessary use of last resort antibiotics and an increased risk of drug resistance ^20^. Recently rapid diagnostic tests have been developed for testing bacterial colonies, utilising antibody mediated capture of carbapenem or ESBL enzymes and providing a visual read out within 15 minutes ^21^. Rapid phenotypic tests based on the colormetric detection of carbapenem hydrolysis are also commercially available, notably the Rapidec Carba NP ^22^, which is sensitive for the detection of most carbapenemases, but insensitive for detecting OXA-type enzymes ^23^.

Molecular assays are also increasingly used to detect resistance markers ^24^. The PCR detection of ESBL and carbapenemase genes has a high correlation with phenotypic testing, and provides faster results 25,26. Molecular testing reduces the time needed to provide targeted therapy ^27^. However, their use as first line diagnostics has been limited because the high number of genes involved necessitates a high degree of multiplexing. For example, a current PCR assay for ESBL alone requires 3 PCR reactions with 7 primer sets, due to their variation across three gene families ^28^. Currently “sample to answer” tests are available such as the GeneXpert CarbaR (Cepheid, US) and FilmArray Blood culture ID (BioFire, US) assays, but their expense makes them difficult to use routinely, and often targeted at high risk sub-populations ^29^. The panels are also not comprehensive, for example the FilmArray assay only includes three resistance markers in its 27 targets, and so provides insufficient evidence to change antibiotic therapy based on likely resistance ^30^.

High resolution melt analysis (HRM) is an endpoint detection method that can be utilised for real time PCR assays to distinguish reaction products based on their melting point. This technique has a greater potential for multiplexing than probe based assays, as it is not limited by the number of channels available in the qPCR system ^31^. HRM assays have been designed for bacterial resistance genes, including carbapenemases ^32–34^, AmpCs ^35^, OXA-48-like beta-lactamases ^36^, chloramphenicol resistance ^37^ and mutations in *aac(6’)-lb-cr* ^38^. However, current assays include a maximum of six targets, and multiple tests are required to thoroughly detect resistance to a drug type. The assay is less costly to operate than probe based approaches, due to the use of an intercalating dye rather than labelled oligonucleotides.

This report describes a two-tube HRM assay for the detection of 14 markers of cephalosporin and carbapenem resistance, including the major carbapenemase, ESBL and plasmid mediated AmpC genes.

## 2. Methods

### 2.1 Assay development

Sequences for each target gene were accessed from GenBank. A minimum of 30 sequences were retrieved for each gene, and care was taken to include a wide coverage of genotypes alongside the reference sequence from the Bacterial Antimicrobial Resistance Reference Gene Database (GenBank Accession PRJNA313047). Sequences of each gene were aligned using Clustal X in MEGA 7, and areas of conservation marked as potential primer binding sites. Primers were designed using Primer3, with a distinct amplicon Tm defined for each target, to enable their differentiation by HRM. A minimum difference of 1°C Tm was selected for each amplicon. The final Tm prediction was obtained using the nearest neighbor calculation on OligoCalc. Primer specificity was checked using Primer BLAST, against the entire non-redundant database. Primers for detecting the carbapenemases KPC, VIM, OXA-48, NDM, and IMP were multiplexed with primers for the three main ESBL groups CTX-M, SHV-like and TEM-like in the ESBL/Carbapenemase assay.

The AmpC assay included the AmpC genes MOX, FOX, EBC, CIT, ACC and DHA, along with a 16S rDNA control. Primers used in each reaction were analysed in all combinations by BLAST to screen for unintended products, and dimer potential was analysed using Multiple Primer Analyzer (ThermoFisher, US).

### 2.2 HRM assays

The ESBL/Carb HRM assay includes 9 primer pairs, and the AmpC assay contains 7 primer pairs including an internal control primer set. Amplicon context sequences are provided in accordance with MIQE guidelines ^39^, in supplementary table 2.

Each HRM reaction contained 12.5 μl Type-it® HRM buffer (Qiagen, UK), primers, and molecular grade water added to a final volume of 20 μl. A 5μl volume of the DNA sample was then added. The qPCR-HRM protocol consisted of an initial denaturation step at 95°C for 5 minutes, followed by 40 cycles of; DNA denaturation at 95°C for 10 seconds, primer annealing at 58°C for 30 seconds, and primer extension at 72°C for 10 seconds with fluorescence monitored in the FAM channel. HRM analysis was carried out over a temperature range of 75°C to 90°C, increased in 0.1°C increments. The results were analysed using the Rotor-Gene Q Software.

The HRM assays were developed using DNA extracted from single isolates in the collection possessing one of the target genes. Due to difficulty in sourcing isolates possessing MOX or ACC genes, plasmids containing the sequences for each gene accessed from GenBank (Accession numbers D13304.2 and AJ133121.1), were synthesised commercially (Oxford genetics, UK). A threshold value for peak calling in the ESBL/Carb assay was set from these initial experiments as 0.4 dF/dT (10% max dF/dT obtained) and retained for all future experiments. The threshold of the AmpC assay was increased to 0.63 to avoid the greater background signal. The amplicon melt temperatures for automatic result calling for each gene are given in Supplementary Table 1, and +/− 0.5C was set as the calling range, except for 16S which was +/− 1C. Due to sequence variation in the CTX-M genes, CTX-M groups one, two and eight have one peak temperature range (denoted group 1), and groups 9 and 25 have another (group 9).

### 2.3 HRM assay evaluation

#### 2.3.1 Strains

A total of 111 well-documented ^40,41^ bacterial isolates from a well characterized collection of clinical isolates sourced in the UK between 2012-2017 were tested. The isolates included *E. coli* (n=37), *K. pneumoniae* (n=27), *Enterobacter cloacae* (n=15), *Enterobacter aerogenes* (n=12), *Citrobacter freundii* (n=12), *Pseudomonas aeruginosa* (n=3), *Morganella morganii* (n=3), *K. oxytoca* (n=1) and *Acinetobacter haemolyticus* (n=1). Isolates were all maintained on Luria-Bertani agar (Oxoid, UK), at 37.5°C. Isolates were phenotypically characterized using the AmpC, ESBL and carbapenemase detection set D72C (Mast Group Ltd, UK), a commercial disk-based method combining cefpodoxime, a penem antibiotic, and various selective beta-lactamase inhibitors, according to the manufacturer’s instructions.

#### 2.3.2 DNA extraction

Genomic DNA was extracted from a single colony of each isolate, suspended in 1ml of sterile water, using a Qiagen DNeasy extraction kit (Qiagen, UK), following the manufacturers protocol for Gram-negative bacteria. DNA was extracted from 200μl of this suspension, and eluted in a volume of 200μl. DNA was stored at −20°C prior to use.

#### 2.3.3 Molecular characterization

All isolates were characterized for ESBL, AmpC and carbapenemase genes using reference assays detailed in ^28^, ^42^ and ^32^, respectively, with some modifications. All thermal cycling parameters were as previously described, DreamTaq PCR reaction mix (Thermo Fisher Scientific, UK) was used for all assays and PCR was performed in the Rotor-Gene Q (Qiagen, UK). Reaction products were analyzed by gel electrophoresis (1% agarose) to confirm expected product size, and select amplicons were bi directionally Sanger sequenced commercially (Source Bioscience, UK) after purification with a QIAQuick PCR purification kit (Qiagen, UK). Reactions were carried out in triplicate, and one or more positive reactions was interpreted as a positive result

#### 2.3.4 Assay evaluation

The extracted DNA from all 111 characterised isolates was tested once with the HRM assays, including the isolates used during the assay development due to the expected low numbers of certain genotypes. In accordance with STARD guidelines ^43^ the HRM assay was carried out by an operator blinded to the reference test results, with positivity called automatically by the RGQ software. The results were then compared against the reference standard PCR and sequencing to estimate diagnostic accuracy.

#### 2.3.5 Intra assay variation

A strain harboring each AMR gene was selected, and the extracted DNA samples were tested in six replicate reactions in a single qPCR run. These results were used to calculate the intra assay variation for each target peak.

## 3. Results

### 3.1 HRM assays

The ESBL/Carbapenemase (ESBL/Carb) HRM assay was able to detect all eight targets, producing a distinct peak for each (Fig.1.A). The mean Tm difference between each peak was 1.5°C (range: 1.9°C –1.1°C), and the minimum difference over 1°C as designed to facilitate differentiation. The AmpC assay generated distinct peaks for each target gene, accompanied by a 16S rRNA control peak in each isolate, apart from the MOX and ACC genes which were synthesised as plasmids (Fig.1.B). The average Tm difference was 1.55°C (range: 1.15° – 2°C). No spurious or unexpected peaks were obtained.

**Figure.1.**
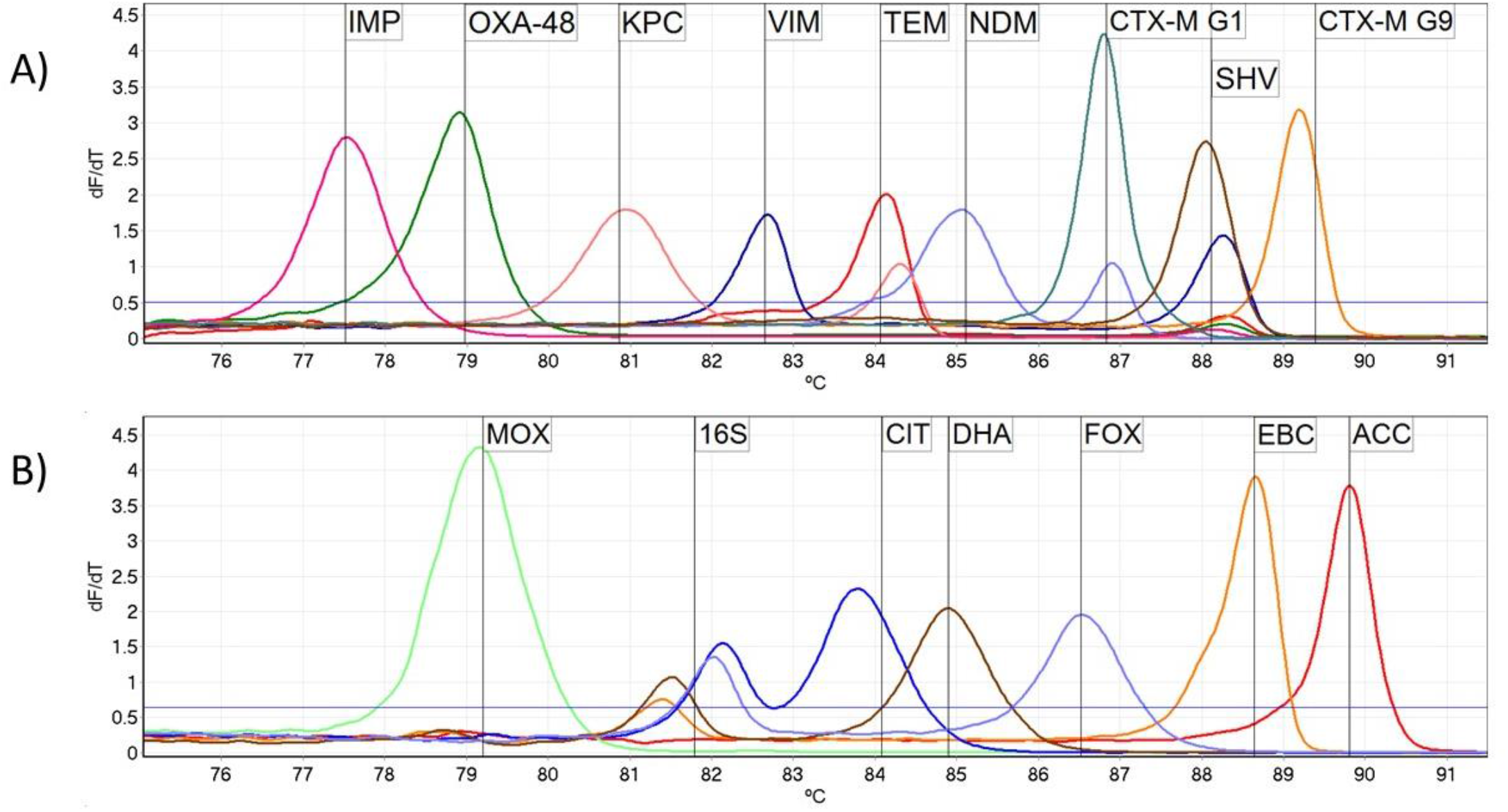
Melt curve profile of the ESBL/Carb assay (A) and AmpC assay (B). The detection threshold is indicated by the solid horizontal line. All isolates in (B) have generated an AmpC peak in addition to a 16S rRNA peak. Two peaks are present in the TEM, CTX-M G1 and SHV bins due to the co-production of these enzymes in isolates used as example carbapenemase producers.

### 3.2 HRM assay evaluation

The assays were evaluated on 111 Gram-negative isolates with mixed resistance patterns. In combination, the two assays correctly identified 97/99 (sensitivity 98%, CI: 92.8%-99.8%) isolates as carrying at least one resistance marker (Table 1.). The sensitivity of the assays for carbapenemase, ESBL and AmpC genes was 96.7% (CI: 82.8-99.9%), 93.6% (CI: 85.7-97.9%) and 93.8% (CI: 82.8-98.7%), respectively. The specificity of the tests for the same genes were 100% (CI: 95.6-100%), 93.9% (CI: 79.8-99.3%) and 93.7% (CI: 84.5-98.2%).

**Table 1.** Diagnostic accuracy of the test for detecting overall resistance and individual genes, with results classified as either TP (true positive), TN (true negative), FP (false positive) or FN (false negative). Results stratified by gene, gene class and presence of any resistance gene (Any R).

| Target | TP | TN | FP | FN | sensitivity (%) | 95% CI (%) | specificity (%) | 95% CI (%) |
| --- | --- | --- | --- | --- | --- | --- | --- | --- |
| **Any R** | 97 | 10 | 2 | 2 | 98.0 | 92.9 - 99.8 | 83.3 | 51.6 - 97.9 |
| **ESBL/Carb** | 79 | 31 | 1 | 0 | 100 | 95.44 - 100 | 96.88 | 83.8 – 99.9 |
| **Carb** | 29 | 81 | 0 | 1 | 96.7 | 82.8 - 99.9 | 100 | 95.6 - 100 |
| **ESBL** | 73 | 31 | 2 | 5 | 93.6 | 85.7 - 97.9 | 93.9 | 79.8 - 99.3 |
| **AmpC** | 45 | 59 | 4 | 3 | 93.8 | 82.8 - 98.7 | 93.7 | 84.5 - 98.2 |
| **IMP** | 1 | 110 | 0 | 0 | 100 | 2.5-100 | 100 | 96.7 100 |
| **VIM** | 4 | 107 | 0 | 0 | 100 | 39.76-100 | 100 | 96.6 100 |
| **NDM** | 6 | 105 | 0 | 0 | 100 | 54.07-100 | 100 | 96.6 - 100 |
| **KPC** | 8 | 103 | 0 | 0 | 100 | 63.1-100 | 100 | 96.48 - 100 |
| **OXA-48** | 10 | 100 | 0 | 1 | 90.9 | 58.7-99.8 | 100 | 96.4 -100 |
| **SHV** | 21 | 76 | 1 | 13 | 61.8 | 43.6 - 77.8 | 98.7 | 93 - 99.9 |
| **TEM** | 37 | 70 | 1 | 3 | 92.5 | 79.6 - 98.4 | 98.6 | 92.4 - 99.9 |
| **CTX-M G1** | 34 | 73 | 2 | 2 | 94.4 | 81.3 - 99.3 | 97.3 | 90.7 - 99.7 |
| **CTX-M G9** | 9 | 102 | 0 | 0 | 100 | 66.4 - 100 | 100 | 96.5 - 100 |
| **ACC** | 0 | 111 | 0 | 0 | - |  | 100 | 96.7 - 100 |
| **CIT** | 15 | 96 | 0 | 0 | 100 | 78.2 - 100 | 100 | 96.2 - 100 |
| **DHA** | 10 | 101 | 0 | 0 | 100 | 69.2 - 100 | 100 | 96.4 - 100 |
| **EBC** | 21 | 83 | 4 | 3 | 87.5 | 67.6 - 97.0 | 95.4 | 88.6 - 98.7 |
| **FOX** | 1 | 110 | 0 | 0 | 100 | 2.5 - 100 | 100 | 96.7 - 100 |
| **MOX** | 0 | 111 | 0 | 0 | - |  | 100 | 96.7 - 100 |
| **16S** | 87 | 0 | 0 | 24 | 78.4 | 69.6 - 85.6 | - |  |

Compared with phenotypic disk diffusion based testing, the HRM assays were 96.7% (CI: 82.8%- 99.9%) sensitive and 100% (CI: 95.5%-100.0%) specific for the detection of carbapenemase resistance. The assay was 88% sensitive (CI: 78.7%- 94.4%) and 94% specific (CI: 80.3% to 99.3%) for detecting 3^rd^ generation cephalosporin resistance (ESBL or AmpC production) in the carbapenem susceptible strains. Of the ESBL/AmpC false negative samples, six were AmpC producers negative for all target genes by reference PCR, and therefore potentially resistant via overexpression of a chromosomal AmpC ^44^.

The ESBL/Carb assay could detect up to three genes simultaneously, including those with neighbouring peaks (Fig.2.A-C). Sensitivity for the particular genes targeted by the ESBL/Carb assay ranged between 90.6 - 100%, with the exception of SHV which was 61.8% (CI: 43.6 - 77.8%) (Table 1). SHV genes were often found in isolates with multiple target genes, and the sensitivity was lower for detecting all genes in co—producers in comparison with isolates with a single gene. Sensitivity in carriers of one, two, three or four genes was 100%, 85.2%, 86.3% and 66.7%, respectively.

**Fig.2.**
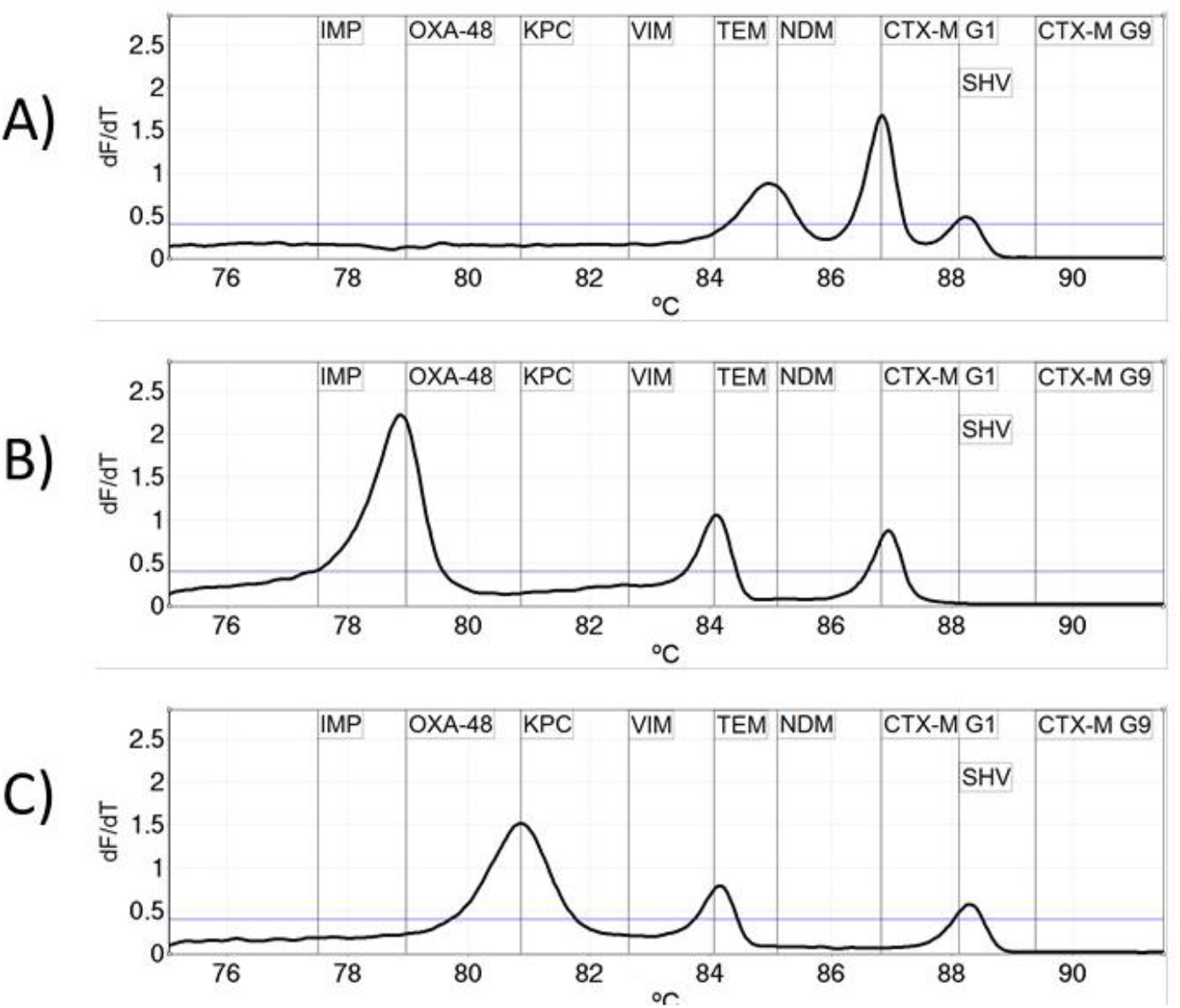
Example melt curves showing the simultaneous detection of three target genes by the ESBL/Carb assay. In all three isolates coexistence of two ESBL genes and a carbapenemase was detected.

The ESBL/Carb assay identified 29/30 carbapenemase producers, failing to identify a single OXA-48 gene. A total of 67/69 isolates (97.1%) were correctly identified as carbapenemase negative but having either ESBL or AmpC genes. There were four false positive results for CTX-M (2) TEM (1) and SHV (1) genes. Sequencing revealed the presence of the intended sequence in all four samples, indicating these samples may have been misidentified by the reference assay.

The AmpC assay identified all AmpC genes with 100% (CI:86.8%-100.0%) sensitivity and specificity, with the exception of EBC genes, which were detected with sensitivity and specificity of 87.5% (CI: 67.6%-97.3%) and 95.4% (88.6%- 98.7%) respectively. The 16S control failed in 24/111 isolates. The concentration of the 16S control primers is deliberately lower than the AMR gene detection sets, to make sure it does not outcompete them in the reaction, and this can lead to false-negative results in isolates with multiple beta-lactamase genes.

The Tm of target genes in both assays all fell within distinct ranges without overlaps (Fig. 3) facilitating discrimination and ensuring the specificity of the HRM assay. The largest range was encountered for the 16S control (81.2-82.33°C), which was expected due to the variability in 16S sequence between bacterial species. This variation was accounted for by the larger calling range for the 16S target, and all 16S peaks were within the selected temperatures. The variation in the AMR targets was markedly reduced, with the smallest range of melt temperatures occurring in DHA (0.12°C), KPC (0.15°C) and OXA-48 (0.2°C). The largest variation was found in the CTX-M group 1 (0.6°C), SHV (0.48°C) and TEM (0.43°C) peaks. The different gene variants included in sample collection are detailed in supplement table 1.

**Fig.3.**
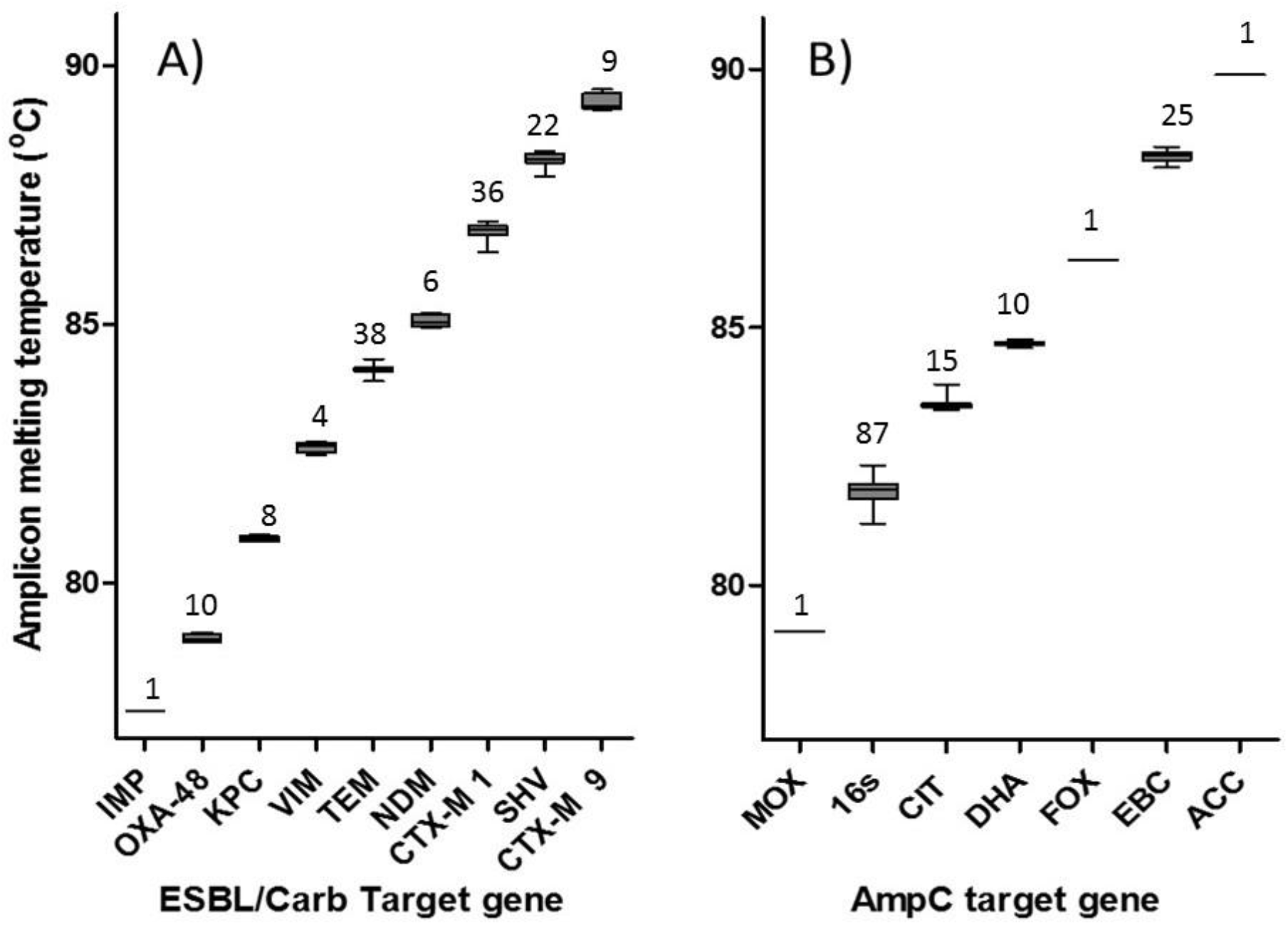
**A)** Tm of the nine amplicons of the ESBL/Carb assay. **B)** Tm of the seven amplicons of the AmpC assay. Numbers indicate total number of peaks represented in the box plots. Whiskers show minimum and maximum Tm.

The intra assay variation for a single producer of each target peak varied between 0.02% and 0.09% in the ESBL/Carb assay, and 0.01% and 0.09% in the AmpC assay (Table 2), indicating a high level of reproducibility between replicates.

**Table 2.**
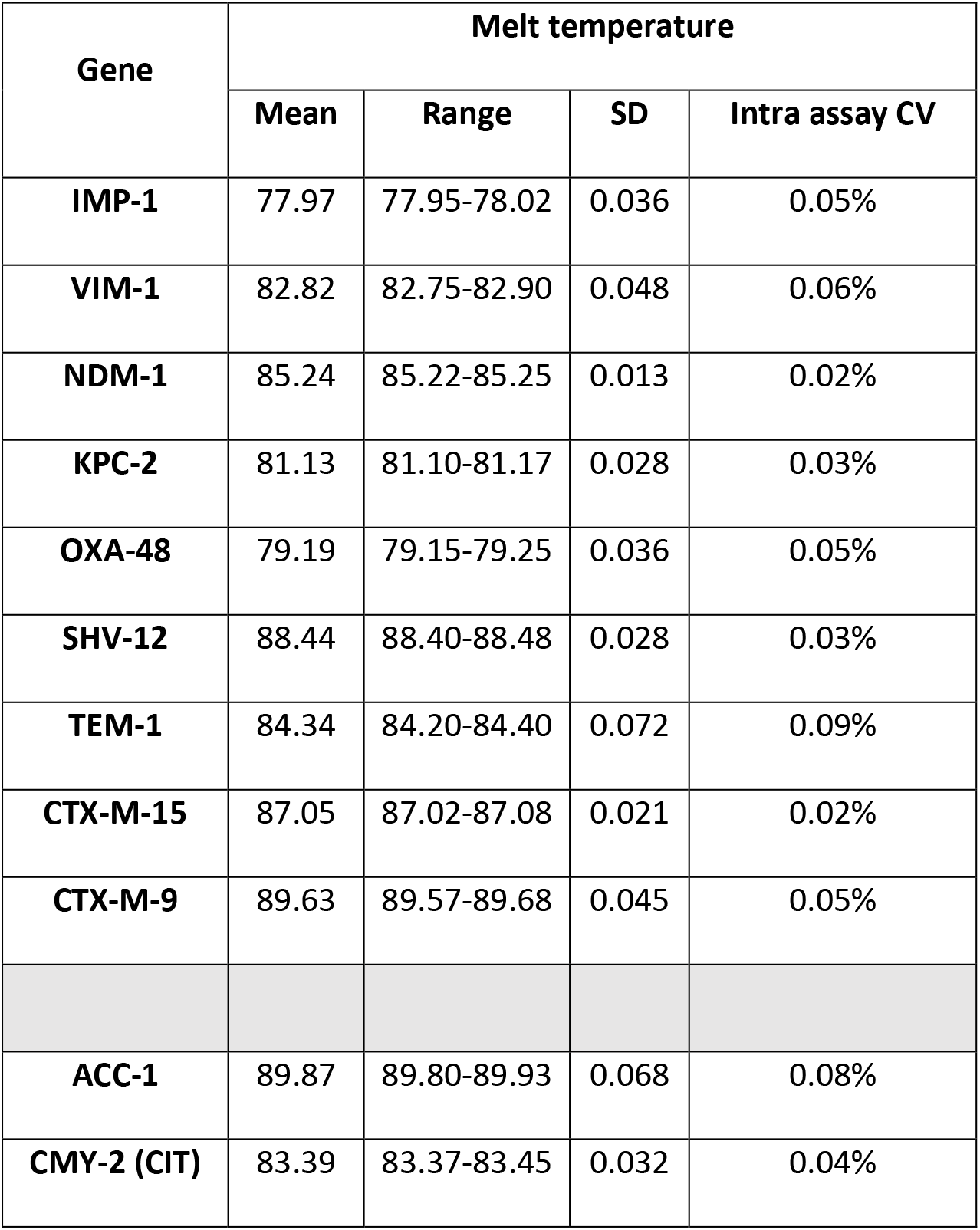
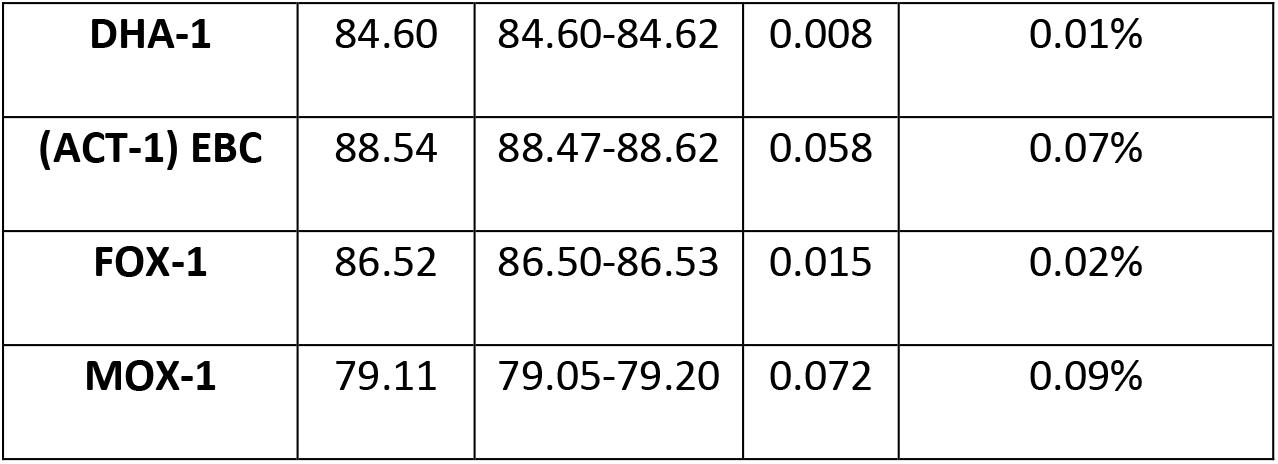
Mean, range, standard deviation (SD) and intra assay coefficient of variation (CV) for six replicate reactions for each target gene

## 4. Discussion

This is the first assay that combines all major ESBL, AmpC and carbapenemase genes into a two-tube assay, enabling the simultaneous detection of 14 of the most important markers of resistance to 3^rd^ generation cephalosporins and carbapenems. Carbapenemase carriage is increasing worldwide, and carriers are typically multidrug resistant, limiting the treatment options and often involving polymyxins or tigecycline ^45^. Resistance to these latter drugs however is also emerging ^46^. Prompt diagnosis would enable the rapid modification of treatment and the instigation of infection control measures to prevent hospital wide outbreaks ^47^.

The ability to simultaneously detect markers of AMR would enable the detection of ESBL and AmpC co-production, which are difficult to detect phenotypically and able to confer resistance to a wider range of antimicrobials including inhibitor/beta-lactam combinations such as amoxicillin/clavulanic acid ^48^.
Screening for multiple genes in this fashion is also useful for surveillance and outbreak monitoring, and gives this test potential utility as an epidemiological tool. The lack of requirement for expensive fluorescent hydrolysis probes, makes HRM a far cheaper way of multiplexing, with an oligonucleotide cost of 3.3% that of a 15 target probe based assay, enabling the detection of a wide range of resistance markers without the costs of sequencing or large qPCR panels.

The assay had high sensitivity of 98% for detecting isolates with any beta-lactamase gene, which is equal to other molecular assays ^26^. There were eight instances across the 111 isolates of genes being identified by the HRM assay that were not identified by the reference standard. The reference tests used were well characterized standard PCRs relying on gel based detection, which can be less sensitive than real-time methods ^49^ and therefore it is possible that these were truly positive isolates. The specificity of HRM assays are typically very high, as they rely on both the specificity of the primer pairs, and the melt temperature of the amplicon generated.

The sensitivity of the test for detecting SHV genes was lower than expected (61%), although the majority of false negative SHV results occurred in isolates with multiple genes. Of the false negative SHV results all the isolates were detected as either ESBL (9/13) or carbapenemase producers (4/13) via different genes and therefore the resistance determination was not affected by not detecting SHV. Whilst co-production of ESBL genes does not affect the susceptibility phenotype of the isolate, and would not impact clinical management of an infection, the ability of the assay to accurately genotype co-producers would be important for its use in surveillance and for improving epidemiological data. Two of the three EBC producers not identified by the assay were *E. cloacae* strains, and the reference PCR may have amplified the chromosomal *E. cloacae* AmpC, which is the ancestor of ACT-1 in the EBC group ^42^. The HRM assay was designed solely using sequence data from plasmid mediated EBC genes, which may explain why it did not successfully identify these targets.

The assay was developed on the Rotor-Gene, but should be transferrable to qPCR platforms designed for HRM, and early experiments on the Magnetic induction cycler (Bio Molecular Systems, Australia) and CFX384 (Bio-Rad, US) have shown equivalent performance.

This initial evaluation was performed using a panel of well characterised and mostly MDR isolates, to ensure a good coverage of target genes. Further evaluations on prospectively collected isolates are required to assess the assay more accurately, including the calculation of predictive values.

The assay detects 14 major carbapenemase, ESBL and plasmid mediated AmpC gene families with sensitivity comparable to other molecular tests, within 3 hours. The assay is highly multiplexed, enabling the comprehensive screening of important beta-lactamases in only two qPCR reactions, which has the potential to save time and costs during testing for these markers. This assay could be used as a cost-effective way to determine the genetic epidemiology of local antimicrobial resistance, or used to provide faster information on antimicrobial resistance from primary cultures than standard phenotypic methods.

## Data availability

All data generated during this study is presented in an analysed format is this manuscript. Raw datasets generated during the current study are available from the corresponding author on reasonable request. Primer mix for the assays are also available for academic partners and can be requested from the corresponding author.

## Author contributions

T.E., C.W., Y.T., J.S., E.R.A., G.H., K.E. and L.E.C. contributed to the conception and design of the study; T.E., Y.T., J.S., S.S., and C.W. carried out all experimental work. T.E., E.R.A. and C.W. analysed the data. T.E. wrote the first draft of the manuscript. All authors were involved in the manuscript preparation and revision, and approval of the final version of the manuscript.

## Acknowledgements

The authors would like to thank the staff at the Liverpool John Moores University School of Pharmacy and Biomolecular Sciences teaching laboratories for assistance with the isolates used in this study.

## Transparency Declaration

All authors declare no conflicts of interest

