## Supplementary tables for "A highly multiplexed melt-curve assay for detecting the most prevalent carbapenemase, ESBL and AmpC genes"

**Institutions**

[1] Research Centre for Drugs and Diagnostics, Liverpool School of Tropical Medicine, Liverpool, United Kingdom.

[2] School of Pharmacy and Biomolecular Sciences, Liverpool John Moores University, Liverpool, United Kingdom

[3]Department of Clinical Sciences, Liverpool School of Tropical Medicine, Liverpool, United Kingdom.

***Corresponding author**

**Running Title:** multiplex melt-curve assay for AMR detection

**Supplementary tables**

**Supplementary table 1.** Gene variants present in the sample collection, and specific gene variants resulting in false negative results in the HRM assay

| GENE FAMILY | GENE VARIANTS | FALSE NEGATIVE |
|---|---|---|
| IMP (1) | IMP-1 (n=1) | |
| VIM (4) | VIM-1 (n-=4) | |
| NDM (6) | NDM-1 (5) NDM-5 (1) | |
| KPC (8) | KPC-1 (1) KPC-2 (2) KPC-3 (4), KPC4 (1) | |
| OXA-48 (11) | OXA-48 (11) | OXA-48 (1) |
| SHV (33) | SHV-1 (22) SHV-12 (4) SHV-18 (1) SHV-27 (5) SHV-110 (1) | SHV-1 (11)<br>SHV-12 (1)<br>SHV-27 (1) |
| TEM (32) | TEM-1 (25), TEM-10 (1), TEM-53 (1), TEM-70 (1), TEM-71 (1), TEM-95 (1), TEM-115 (1) TEM-214 (1) | TEM-1 (3) |
| CTX-M G1 (36) | CTX-M-1 (2), CTX-M-3 (6) CTX-M-15 (22) CTX-M-28 (6) | CTX-M-15 (2) |
| CTX-M G9 (9) | CTX-M-9 (7), CTX-M-14 (1) CTX-M-27 (1) | |
| ACC (0) | ACC-1 plasmid | |
| CIT (14) | CMY-2 (10) LAT-4 (1), CMY-8 (1), CMY-86 (1), CMY-112 (1) | |
| DHA (10) | DHA-1 (8) DHA-2 (2) | |
| EBC (25) | ACT-1 (12) ACT-17 (2) ACT-18 (1), ACT-23 (1), ACT-30 (1), ACT-31 (2), ACT-32 (4), ACT-36 (2) | ACT-1 (3) |
| FOX (1) | FOX-1 (1) | |
| MOX (0) | MOX-1 plasmid | |

**Supplementary Table 2**. Context sequence for the amplicons form the ESBL/Carb and AmpC assay, provided in accordance with MIQE guidelines. The amplicon context sequence includes the amplicon location, with 20bp unevenly distributed to each flanking region.

| Assay | Target Gene | GenBank accession | Amplicon context sequence | Amplicon size (bp) | Amplicon Tm (°C) |
|---|---|---|---|---|---|
| ESBL/Carb | TEM | NG_050145.1 | 581-805 | 204 | 84.2 |
| | SHV | NG_049989.1 | 20-239 | 199 | 88.2 |
| | CTX-M Group 1, 2, 8 | NG_048935.1 | 613-863 | 230 | 86.8 |
| | CTX-M Group 9, 25 | | | | 89.4 |
| | IMP | NG_049172.1 | 207-327 | 100 | 77.7 |
| | OXA-48 | NG_049762.1 | 226-376 | 130 | 79 |
| | KPC | NG_049253.1 | 258-366 | 88 | 81 |
| | VIM | NG_050336.1 | 285-455 | 150 | 82.7 |
| | NDM | NG_049326.1 | 370-491 | 101 | 85.3 |
| AmpC | CIT (CMY- 2, 4, 6, 7, LAT- 1- 4, BIL- 1) | NG_048814.1 | 290-420 | 110 | 83.52 |
| | DHA (DHA- 1, 23, 24) | NG_049055.1 | 105-238 | 113 | 85.1 |
| | EBC (ACT- 3, 5, 14, 18, 21, 23, 24, 36, 39, MIR- 1, 3, 5- 8, 11, 12, 17, 18) | NG_049293.1 | 725-1144 | 399 | 88.55 |
| | FOX (FOX- 1- 9, 12, 13) | NG_049098.1 | 400-520 | 100 | 86.45 |
| | MOX | NG_049315.1 | 93-431 | 318 | 89.36 |
| | ACC | NG_048588.1 | 407-538 | 111 | 77.3 |
| | 16S | CP028994.1 | 1545009-1545241 | 212 | 84.48 - 84.52, |
